# Sparse Coding with a Somato-Dendritic Rule

**DOI:** 10.1101/451955

**Authors:** Damien Drix, Verena V. Hafner, Michael Schmuker

## Abstract

Cortical neurons are silent most of the time. This sparse activity is energy efficient, and the resulting neural code has favourable properties for associative learning. Most neural models of sparse coding use some form of homeostasis to ensure that each neuron fires infrequently. But homeostatic plasticity acting on a fast timescale may not be biologically plausible, and could lead to catastrophic forgetting in embodied agents that learn continuously. We set out to explore whether inhibitory plasticity could play that role instead, regulating both the population sparseness and the average firing rates. We put the idea to the test in a hybrid network where rate-based dendritic compartments integrate the feedforward input, while spiking somas compete through recurrent inhibition. A somato-dendritic learning rule allows somatic inhibition to modulate nonlinear Hebbian learning in the dendrites. Trained on MNIST digits and natural images, the network discovers independent components that form a sparse encoding of the input and support linear decoding. These findings confirm that intrinsic plasticity is not strictly required for regulating sparseness: inhibitory plasticity can have the same effect, although that mechanism comes with its own stability-plasticity dilemma. Going beyond point neuron models, the network illustrates how a learning rule can make use of dendrites and compartmentalised inputs; it also suggests a functional interpretation for clustered somatic inhibition in cortical neurons.

## Introduction

Activity in the brain is sparse: a given pyramidal neuron spikes infrequently (lifetime sparseness), and few neurons are active at once (population sparseness). This coding scheme is efficient in terms of energy use, and also in terms of storage capacity [1]: the same cell can participate in multiple, overlapping assemblies [2] that are active in different contexts.

Beyond energy and storage efficiency, the promise of sparse codes is that they can reveal structure in natural inputs which makes it easier to learn associative mappings: detect a stimulus, transform a pattern of neural activity into motor commands, or prime the activation of another cell assembly. In some sense, every pathway that links two populations of neurons involves a transformation of one neural code into another, and a sparse, decorrelated code can reduce the computational cost of these transformations by allowing linear readouts [3,4].

The classical way to learn sparse codes involves the information-maximization principle of Bell & Sejnowski [5] for blind source separation. In independent component analysis [1,6,7] and sparse auto-encoders [8], the algorithm works to minimise a global cost function that includes a sparse constraint. Here, we focus on a family of single-layer networks that do not compute a global cost explicitely. Instead, these networks learn sparse codes with local learning rules thanks to the combination of two unsupervised heuristics: projection pursuit and competitive learning.

Projection pursuit looks for receptive fields with a non-Gaussian activity distribution. Diaconis & Freedman [9] note that these tend not to occur by chance, reflecting instead some fundamental structure in the input — a characteristic that reminds us of the *suspicious coincidences* of Barlow [10].

As for competitive learning, described by Rumelhart & Zipser [11], it aims to reduce the redundancy of the code and decorrelate the output dimensions, so that each neuron responds to a different feature. This usually involves a winner-take-all system [12], or inhibitory connections between the coding neurons [13,14] — an organisation which is equivalently called lateral, recurrent or mutual inhibition.

Starting with Földiák [15], these two heuristics have been applied in a variety of sparse coding networks with rate-based [16–18] and then spiking neurons [19–22]. These networks have in common the use of Hebbian lateral inhibition to decorrelate the output, and of nonlinear Hebbian rules to perform projection pursuit on the feedforward input.

Nonlinear Hebbian learning, to follow the terminology of Brito & Gerstner [23], refers to a variant of Hebbian learning where the change of weight is proportional to the correlation between the input and a nonlinear function of the output (more precisely, of the receptive field activation). The Bienenstock–Cooper– Munro (BCM) rule [24] is an early example of such a rule, inducing depression when the output activity is below average and potentiation when it is above average. This steers gradient descent towards an activity distribution with heavy tails, which typically converges onto one of the independent components.

As noted by Brito & Gerstner [23], the precise shape of that nonlinear function is not critical. The trick is to keep it aligned with the activity distribution throughout learning, so that the potentiation region stays centered on the tail. Usually, this is done by enforcing a constant norm for the weight vectors, or by using a homeostatic term that moves the potentiation threshold according to the average activity of the neuron, as in the BCM rule. That homeostatic term has the effect to regulate the lifetime sparseness of the neuron and is also called *intrinsic plasticity* (IP) by Triesch [25], to distinguish it from synaptic plasticity.

In most models, IP needs to be faster than the Hebbian component of learning [26,27]. But in vivo, IP tends to be slower, acting over a timescale of days rather than minutes [29,30]. Besides, fast firing rate homeostasis could be particularly disruptive for animals and robots that learn continuously, and cannot assume that the feature detectors they have acquired will be stimulated at regular intervals.

Here we propose an alternative scheme that does not require fast intrinsic plasticity. The idea is to put mutual inhibition itself in control of the Hebbian nonlinearity: stimuli for which many neurons compete to respond, and neurons that are often active as well, would attract more lateral inhibition and be subject to a higher potentiation threshold. In other words, instead of using intrinsic plasticity to enforce lifetime sparseness, this scheme would regulate both the population and the lifetime sparseness through synaptic plasticity.

To do so, we need a mechanism through which the feedforward learning rule could measure the amount of competition on an input-by-input basis and use it as a negative feedback. But artificial neural networks usually employ point neurons, where all inputs are added together into a single activity variable. The consequence is that the learning rule cannot distinguish between stronger lateral inhibition — the signal to become more selective — and weaker feedforward activity that results from synaptic plasticity or from fluctuations in the input.

The solution could be to integrate the feedforward and recurrent pathways in separate neural compartments, for instance the soma and a dendrite. The dendritic compartment could then estimate the amount of somatic inhibition by comparing its local depolarisation with the somatic activity that it perceives via backpropagating action potentials.

The idea has been tried before, although not on a sparse coding task. In Körding & König [31], lateral inhibition can prevent the backpropagating action potentials from reaching the dendrites, which induces depression in dendritic synapses via spike-timing dependent plasticity. Urbanczik & Senn [32] use probabilistic spiking neurons where the dendritic compartment tries to match the somatic potential; this results in depression when unpredicted external inputs inhibit the soma, and potentiation when these unpredicted inputs are excitatory instead.

Here we set out to investigate whether a variant of these somato-dendritic learning rules could discover sparse codes in natural stimuli. We found that one can adjust the somatic and dendritic transfer functions to produce a BCM-like curve where the threshold between depression and potentiation follows an instantaneous measure of somatic inhibition. This lets the network learn sparse codes by regulating population sparseness instead of lifetime sparseness, and does not require fast intrinsic plasticity.

## Results

### Network model

Our model is a network of neurons, each of which consists of a spiking, leaky integrate-and-fire (LIF) soma, and a rate-based dendritic compartment (fig. 1). We summarise its main features here and refer the reader to the *Methods* section for the full details.

The dendrites have distinct receptive fields *w* and integrate feedforward input rates *x* for each stimulus, yielding a current *I*_*d*_:

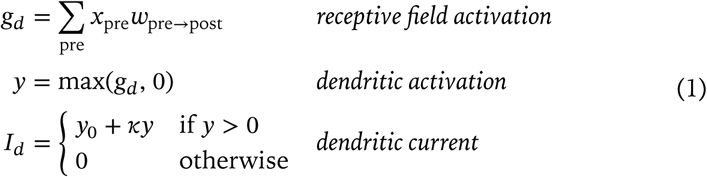

The somas integrate both the dendritic current *I*_*d*_ and a current *I*_*s*_ from recurrent inhibitory synapses to drive a varying membrane potential *u*:

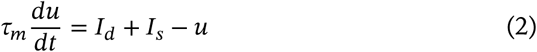

Somas emit spikes according to standard LIF dynamics with a fixed threshold, fixed reset and no refractory period. These spikes induce recurrent inhibition throughout the network via the current *I*_*s*_, and are also used to compute a firing rate *z* that modulates learning in both the dendritic and the somatic synapses.

The network is meant to model a small patch of neural tissue where full connectivity is an acceptable approximation; hence we keep the number of neurons small (*N* ≤ 1024). With respect to the dimensionality *d* of our input stimuli, this translates to networks that range from undercomplete (*N*/*d* ≪ 1) to slightly overcomplete (*N*/*d* ≈ 1.3).

**Figure 1:**
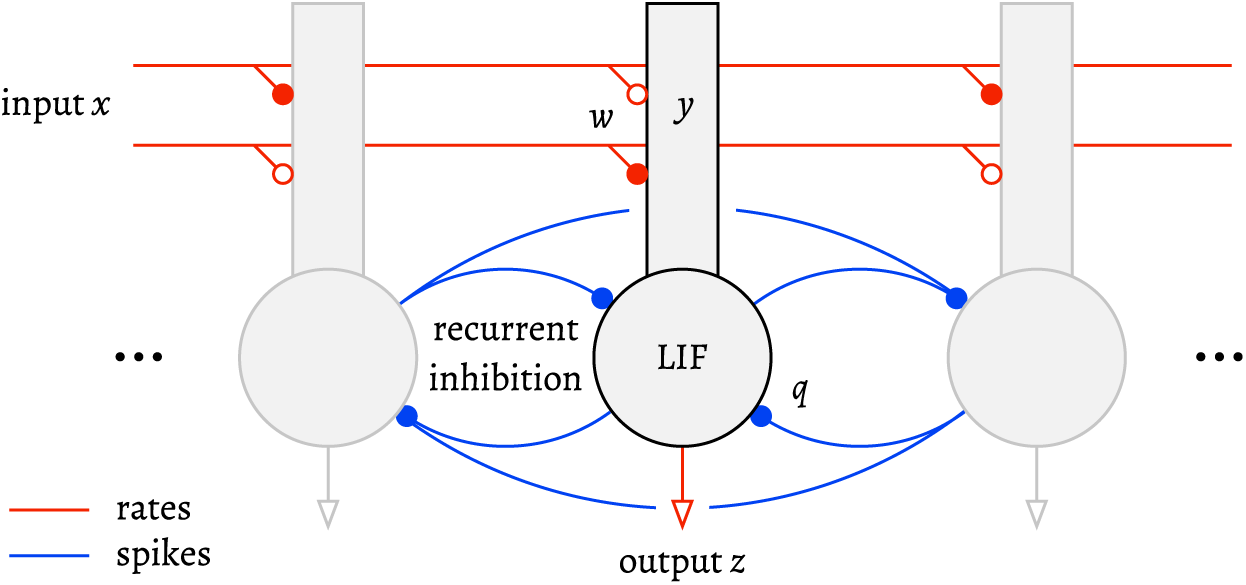
Architecture of the network. Annotations indicate the feedforward input *x*, leaky integrate-and-fire (LIF) somas and their firing rate *z*, dendritic compartments and dendritic activity *y*, and feedforward and recurrent pathways with weights *w* and *q*, respectively. The symbol • denotes an inhibitory synapse, ∘ an excitatory one.

There are two fully-connected pathways: a recurrent inhibitory pathway between the somas, and a feedforward pathway between the input and the dendrites.

The feedforward pathway targets the dendrites and contains both excitatory and inhibitory synapses. It carries rates instead of spikes; doing so allows us to employ a classical Hebbian formalism in the learning rule and discrete-time dendritic compartments. A spike-based input and continuous-time dendrites would be more biologically plausible, but the model would also become substantially more complex; we reserve these for future work. Here we use a rectified linear activation function in the dendrites, with some modifications to account for the overal transfer properties of biological dendrites (see *Methods* for details).

The recurrent pathway mediates all-to-all inhibition via spikes and conductance-based somatic synapses. For simplicity we do not use separate inhibitory interneurons (although that architecture deviates from biology and Dale’s law, King et al. [21] found that replacing direct inhibition with interneurons did not substantially alter the results of Zylberberg et al. [20]). We do not model propagation delays which are de-facto fixed at one timestep *dt*. We allow self-inhibition for simplicity, as it has only a minor effect on receptive field formation (fig. 5). Self-inhibition decreases the slope of the current-frequency (I-ƒ) curve of the LIF neuron without changing its threshold (it acts like a relative refractory period and can only affect future spikes). Thus it can in theory be compensated for by parameters controlling the input gain.

The network operates as follows. We present each input pattern *x* to the dendrites and compute the dendritic activation *y*. This results in a constant current flow *I*_*d*_ from the dendrite to the soma while the somas compete to respond for 100 timesteps (*dt* = 0.5 ms), producing spikes that induce time-varying inhibitory currents *I*_*s*_ Then we compute firing rates *z* using both the number of spikes and the spike latencies. Finally, we apply the feedforward and recurrent learning rules. We repeat these steps for the next input pattern, etc.

### Feedforward learning rules

The weight *w* of each feedforward, dendritic synapse is updated according to a nonlinear Hebbian rule:

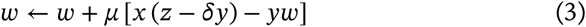

where *x* is the input rate, *y* is the dendritic activation, *z* is the somatic firing rate, *µ* is the learning rate and *δ* sets the potentiation/depression ratio. The rule can change the sign of the weights, switching between excitatory and inhibitory synapses.

The core of the learning rule is the term *z* − *δ*_*y*_ (fig. 2). Within that term, *y* is non-linear with respect to the receptive field activation *g*_*d*_ (due to the dendritic rectification), and *z* is itself nonlinear with respect to *y* (due to the somatic response threshold, which increases with somatic inhibition). Thus *z* −*δy* is zero for sub-threshold inputs (*gd* ≤ 0 and *z* = *y* = 0), negative for super-threshold inputs but weak somatic responses (*gd* > 0 and *y* > 0, but *z* < *δy*), and positive for strong somatic responses (*gd* > 0, *y* > 0, and *z* > *δy*).

This mirrors the term *y* (*y* − 〈*y*^2^〉) in the BCM rule, which is also non-linear with respect to the receptive field activation, inducing long-term depression (LTD) for weak responses and long-term potentiation (LTP) for strong responses. But where the BCM rule defines weak and strong responses in relation to the average activity 〈*y*^2^〉 (lifetime sparseness), here we define them in terms of winning or losing the competition to respond (population sparseness).

If the dendrite is active (*y* is large) but the soma is inhibited (*z* is comparatively small), the rule induces LTD: the neuron tried to respond and lost to more active neurons. If the dendrite is active and the soma responds strongly (*z* > *δy*), the rule induces LTP: the neuron is one of the winners. If the dendrite is not active (*y* = 0), the soma is not active either (*z* = 0) and there is no synaptic change: the neuron did not participate in the competition for that particular input.

**Figure 2:**
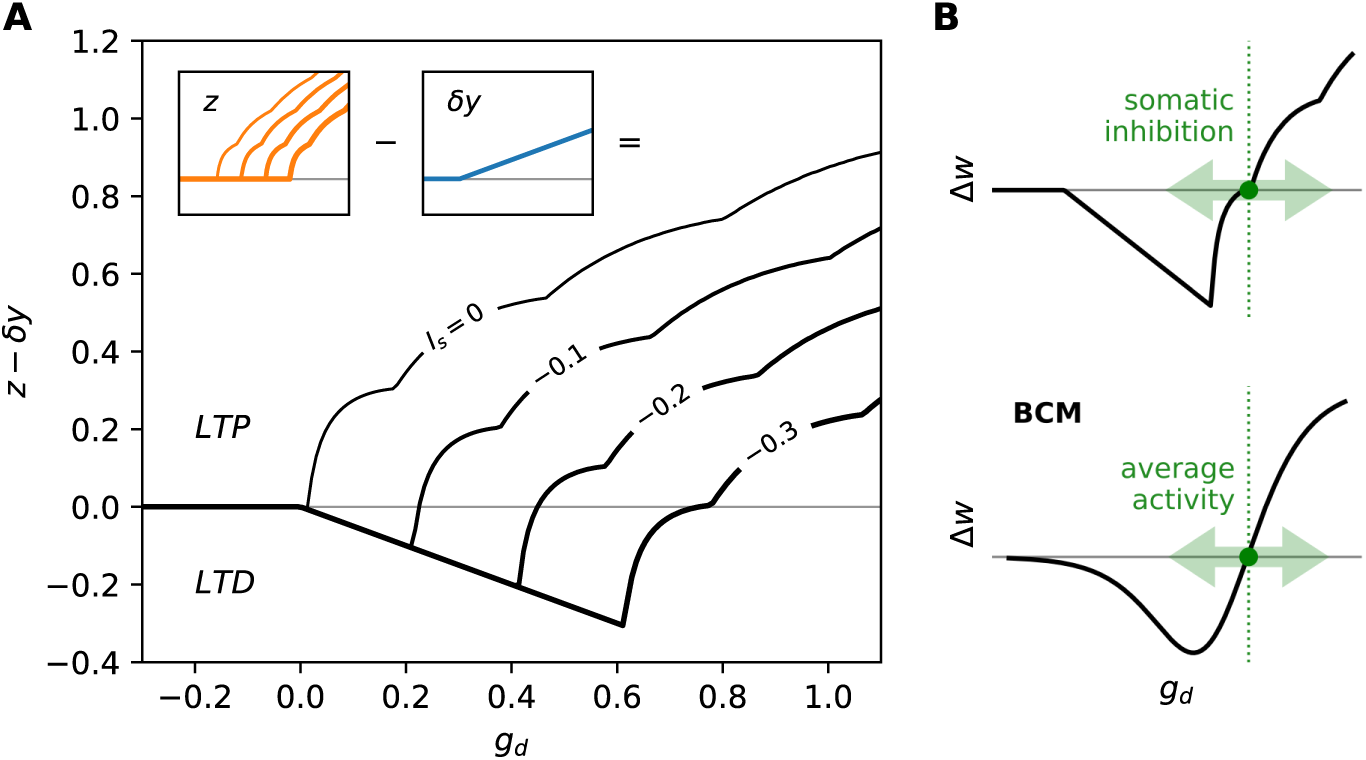
The learning rule produces a BCM-like nonlinearity controlled by somatic inhibition. **A:** effective Hebbian nonlinearity *z* − *δ*_*y*_ as a function of the receptive field activation *g*_*d*_ and somatic inhibition. Injecting a constant inhibitory current *I*_*s*_ into the soma (marked on the curves) shifts the potentiation threshold to the right. The contributions of the terms *y* and *z* are shown in insets. Bumps in the curves are a consequence of the way we compute the firing rate *z* and mark the occurence of an extra spike. **B:** the result is a BCM-like nonlinearity controlled by somatic inhibition, whereas the BCM rule itself is controlled by average activity. *Note: this figure was generated with a finer timestep* dt = 0.01 ms *to smoothe the discontinuities in the curves caused by the discrete spike times.*

The decay term −*yw* sets a steady-state value for the weights and scales the maximum dendritic activity *y* as a function of the receptive field size. It is gated by dendritic activity: there is no decay when *y* = 0. This ensures that the weights do not fade when the dendrite is inactive.

Finally, we apply a separate regularisation rule after the Hebbian changes, taking care not to change the sign of the weight:

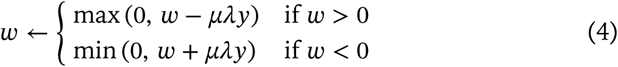

where λ determines the amount of regularisation. This does not fundamentally change the operation of the learning rule, but simplifies the receptive fields by suppressing the weights of weakly correlated input dimensions.

In summary, the feedforward learning rule compares the dendritic and somatic activity to estimate whether the neuron was silent, losing or winning. It then updates the weights so that inputs correlated with losing become inhibitory, inputs correlated with winning become excitatory, and inputs correlated with being silent are removed from the neuron’s receptive field.

### Recurrent learning rule

The somatic synapses that mediate lateral inhibition are plastic as well. The weight *q* of each recurrent, somatic synapse between a pre- and a post-synaptic neuron follows a standard Hebbian rule with pre-synaptic gating:

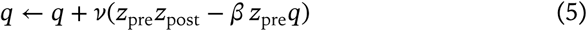

where a constant υ sets the learning rate and another constant *β* controls the scale of the weights (see *Methods* for parameter values). Gating by *z*_pre_ ensures that the inhibition from a winning neuron to a losing neuron decays, but the reciprocal connection does not. The asymmetry prevents a single neuron from taking over all the input features [33]. In practice, we use a much faster learning rate for the recurrent inhibition compared to the feedforward synapses (*ν* ≫ *µ*); otherwise receptive fields are unstable and oscillate between selective and nonselective features.

Inhibitory plasticity, as opposed to fixed inhibition, has two roles in our model. First, it ensures that neurons compete with each other only to the degree that their responses are correlated [33]. Thus if two neurons respond to features that are only weakly correlated, they can occasionaly be strongly active at the same time without influencing each other. And second, it stabilises the feedforward learning rule just like the homeostatic threshold does in the BCM rule: it ensures that as neurons fire more they also attract more inhibition, which prevents the distribution of *g*_*d*_ from escaping the LTD region of the feedforward learning rule (fig. 2). If the recurrent inhibitory weights were fixed, all dendrites would learn the same non-selective receptive field (fig. 5).

### Receptive fields

Our first experiment is to look at the receptive fields of the neurons after training on various types of inputs. The expectation, for a sparse coding network, is that these receptive fields should correspond to selective features (rather than whole input patterns) and that the neurons should be silent most of the time.

Trained on the MNIST dataset of handwritten digits [34], the network learns receptive fields that respond to fragments of digits or pen strokes, as shown in fig. 3. These receptive fields resemble the ones learned by sparse auto-encoders [8], despite the fact that we use a different algorithm — a coincidence which can be explained if these pen-stroke shapes are indeed the independent components of MNIST digits. We also test a variant of MNIST called Fashion-MNIST [35], which uses the same format but consists of small images of items of clothing like shoes and shirts. Training the network on that dataset extracts the outlines of the input stimuli and also separates some of their constituent parts.

**Figure 3:**
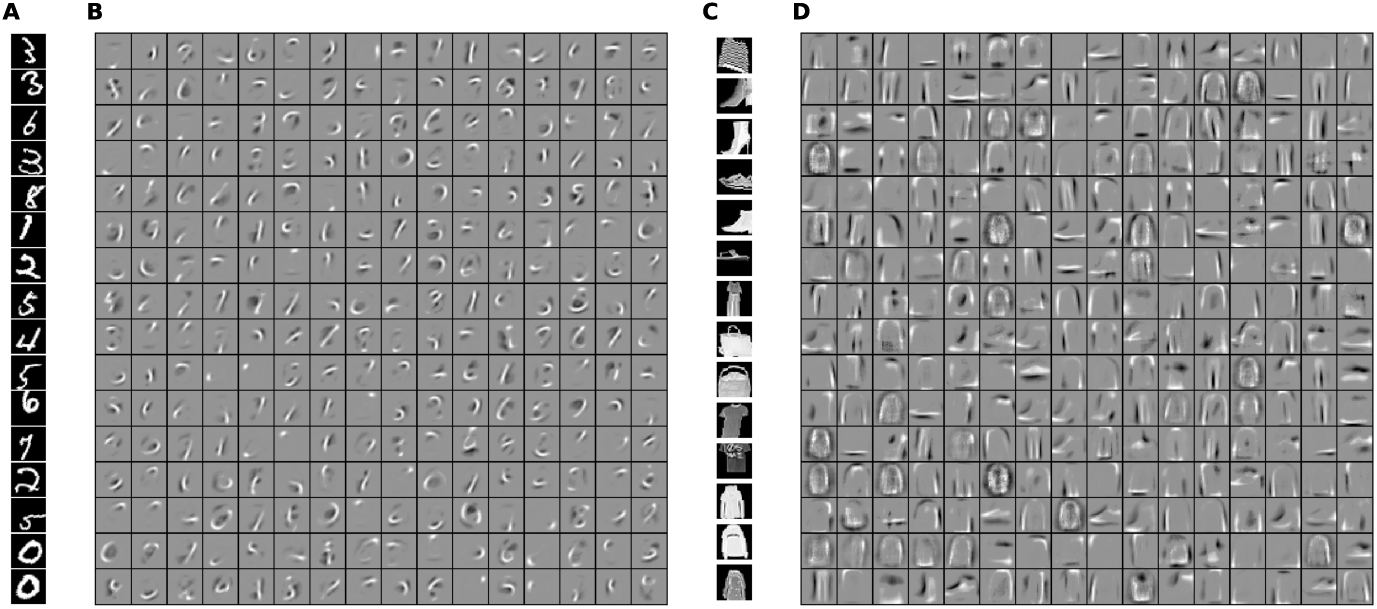
The network learns independent components from the MNIST datasets. **A, B**: the network learns pen-stroke shapes from the MNIST dataset. **A**: sample input stimuli. Black corresponds to zero and white to one. **B**: receptive fields (weights) of a network with 256 neurons after training on 120,000 digits (28 × 28 pixels) with random distortions. Middle gray corresponds to zero, lighter pixels to excitatory weights, and darker pixels to inhibitory weights. **C, D**: the network learns the outlines and parts of the various items of clothing in the Fashion-MNIST dataset; for instance the neuron in the top-right corner of **D** responds to short sleeves. All other details are the same as for **A** and **B**.

We then train the network on two photographic datasets. The first one is the dataset used by Olshausen & Field [1], which consists of landscapes and closeups of natural outdoors scenes. The second one, which we refer to as the Monuments dataset, is a selection of black-and-white archive photographs from the Cornell University Digital Collections [36] that show monuments and cities of France. Sparse coding networks have often been applied to natural images [1,17,19,20], from which they learn Gabor-like filters that resemble the receptive fields of simple cells in the visual cortex [37]. Images are typically not presented to the network in their raw form, but first processed either by a *differenceof-Gaussians* filter that models the transformations happening in the retina, or by a whitening transform that equalises the variance across spatial frequencies [38]. Both types of pre-processing have the effect to suppress low spatial frequencies and highlight edges. For this experiment we adopt the whitening transform of Olshausen & Field [1].

After training on small patches drawn at random from different image locations, the model learns oriented edge filters, in line with other sparse coding algorithms (fig. 4). Compared to the outdoor scenes used by Olshausen & Field [1], the Monuments dataset yields more elongated receptive fields; this is probably due to the more frequent occurence of straight edges in scenes that contain man-made objects.

**Figure 4:**
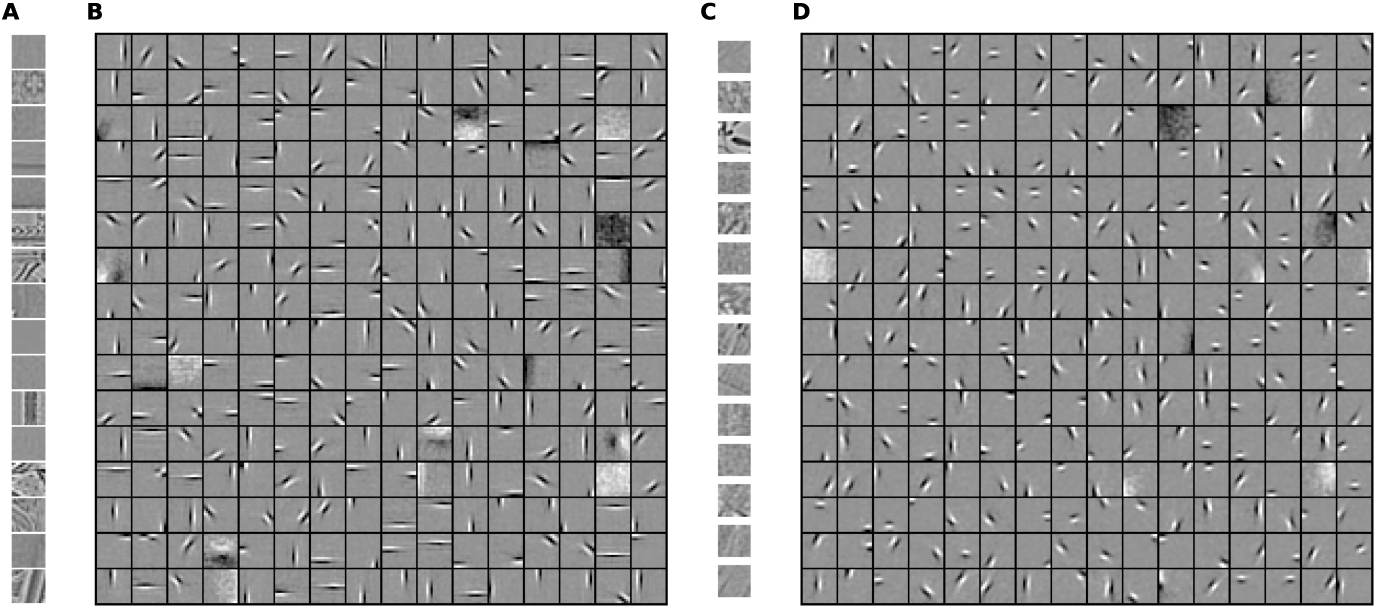
The model learns oriented edge filters from pre-whitened natural images. **A, B**: Monuments dataset. **A**: sample input patches. Middle gray corresponds to zero. **B**: receptive fields after training on 500,000 patches (16 × 16 pixels) (see *Methods*). In addition to the localised edge filters, the network develops a pair of non-selective ON and OFF receptive fields (uniform dark and bright patches). These encode the mean of the input, which we do not subtract. Similarly, a dozen cells learn broad oriented gradients. **C, D**: Olshausen & Field dataset. Receptive fields tend to be shorter and more localized than with the Monuments dataset. All other details are the same as for **A** and **B**.

### Linear decoding

The next series of experiments aims to check whether the network’s output is indeed a good encoding of the input. This does not necessarily follow from an analysis of the receptive fields; for instance, a network could succeed in extracting individual independent components, but still fail to encode the mixture of components present in any given input. More specifically, we would like to check whether the sparse encoding produced by the network can be linearly decoded, enabling cheap multiple readouts of cell assemblies as envisioned by Fusi et al. [4].

**Figure 5:**
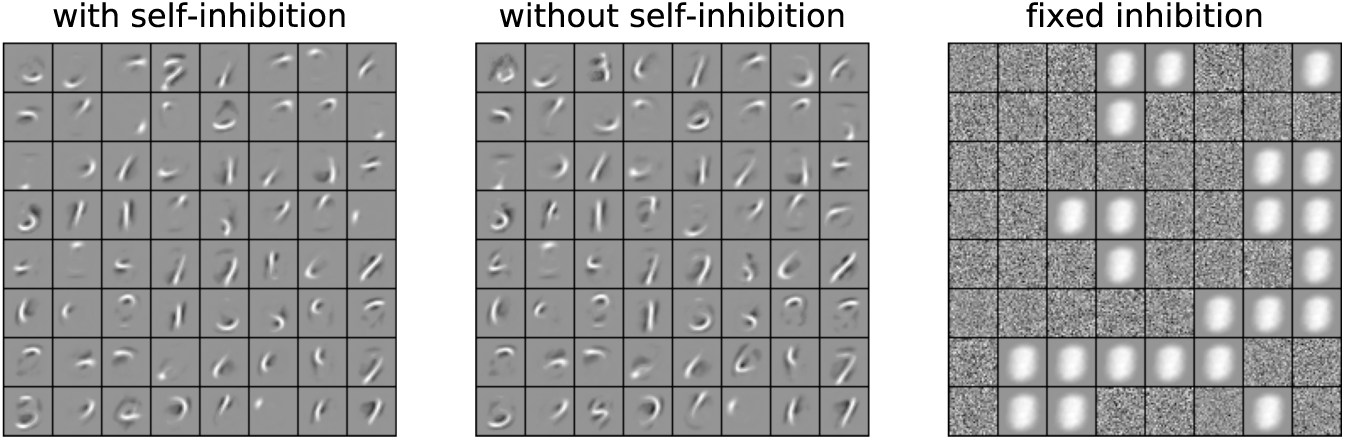
Inhibitory plasticity, but not self-inhibition, is required for the learning rule to function. Receptive fields from three networks with *N* = 64 neurons after learning from the same initial random state, but with different configurations of recurrent inhibition. In the variant with fixed inhibition, some neurons stop responding early in the learning process and still have a random receptive field; all active neurons have the same receptive field which corresponds to the average digit.

We first test whether sparse codes make it easier to classify MNIST digits (tbl. 1). Trained on the raw pixels, a linear Support Vector Machine (SVM) classifier performs poorly on MNIST, with an error rate of 8.2 %. But the same linear classifier reaches a much better performance if we train it on the output of the sparse coding network instead of the raw pixels. With *N* = 512 neurons, that combination outperforms the non-parametric k-Nearest Neighbors method (kNN). It also compares with a Multi-Layer Perceptron (MLP) with three layers — in that particular case, the unsupervised sparse coding layer effectively replaces two hidden layers trained using backpropagation. With *N* = 1024 neurons, the accuracy reaches a value of 0.981 that is higher than most non-convolutional methods with the exception of polynomial SVM [35,39].

**Table 1:** The network learns features that improve linear decoding. Mean and standard deviation of the test error on MNIST (10 runs with different random seed). For comparison we include data for a MLP (28×28–300–100–10) from LeCun et al. [39]; other results are our own work.

| Input | Classifier | Error (%) |  |
| --- | --- | --- | --- |
| Raw pixels | SVM (linear) | 8.2 |  |
| Raw pixels | kNN ($k = 4$) | 3.2 | |
| Raw pixels | MLP (3 layers) | $2.5 \pm 0.1$ [39] | |
| Sparse ($N = 256$) | SVM (linear) | $3.3 \pm 0.2$ | |
| Sparse ($N = 512$) | SVM (linear) | $2.3 \pm 0.1$ | |
| Sparse ($N = 768$) | SVM (linear) | $2.1 \pm 0.1$ | |
| Sparse ($N = 1024$) | SVM (linear) | $1.9 \pm 0.1$ | |

We obtain broadly similar results with the Fashion-MNIST variant (tbl. 2): sparse coding improves linear decoding. However, this dataset is more challenging than the standard MNIST digits for non-convolutional algorithms, and the improvement is consequently smaller. Ensemble learning methods as well as polynomial SVM yield a better performance [35].

**Table 2:**
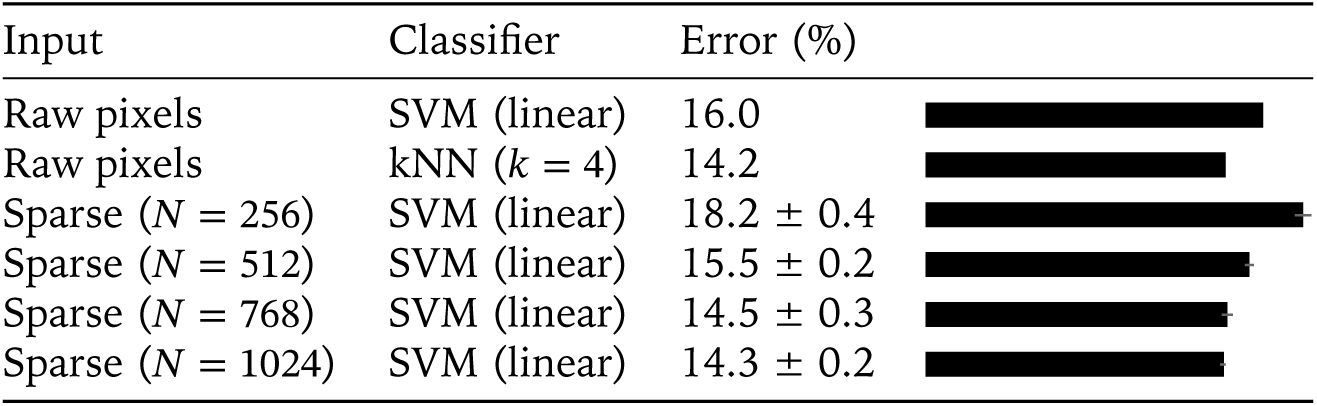
Sparse features also improve linear decoding on the FashionMNIST dataset, although less than with the standard MNIST.

After that classification task, we turn to linear regression and attempt to reconstruct natural images from the output of the network. While Zylberberg et al. [20] inverted the transformation manually by reusing the network’s encoding weights for decoding, here we train a linear model to predict the input patch given the sparse output of the network. We did not attempt to quantify the reconstruction error: pixel-wise measures such as the peak signal-to-noise ratio are neither very informative of how much structure is preserved, nor easy to interpret when comparing different scenes, and better metrics based on structural similarity are non-trivial to compute [40]. Qualitatively, we find that even a small network with 64 neurons preserves the general features of the scene (fig. 6), despite reducing the dimensionality of the data by a factor of 4.

**Figure 6:**
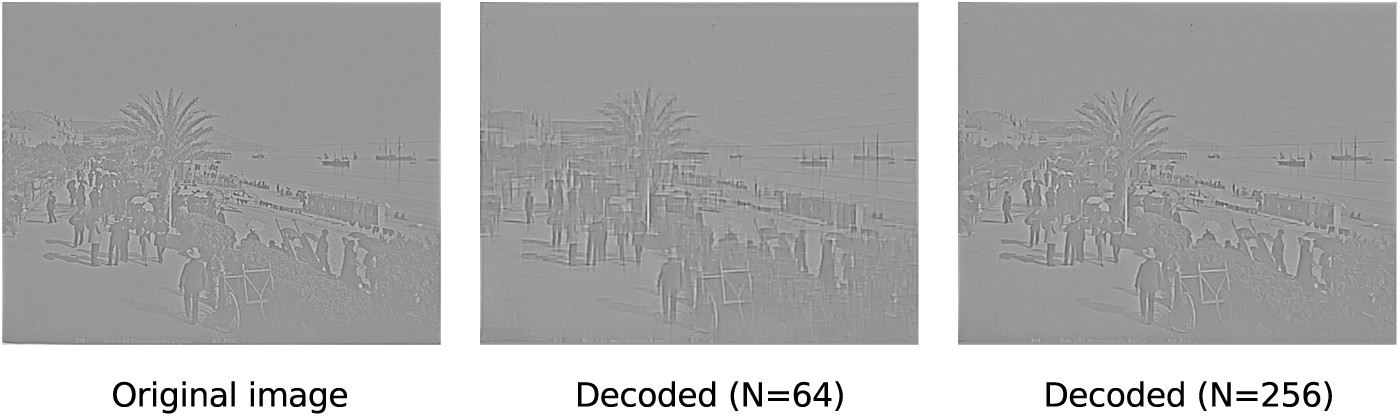
Input images can be linearly decoded from the sparse output. Reconstructions from networks of 64 to 256 neurons show increasing fidelity to the original image.

### Sparseness

The activity of the network is sparse at the end of the training period, both in terms of lifetime and population sparseness (fig. 7). Plastic recurrent inhibition rapidly enforces sparse spiking and maintains it at the level of a Poisson process with the same rate, or slightly higher (fig. 8).

Lehky et al. [41] make the point that lifetime and population sparseness in sensory neurons are inter-related: if the responses of the neurons are uncorrelated, then their population and lifetime sparseness must be equal, a property they call ergodicity. We find that this is indeed true of our network: for all the datasets we tested, both types of sparseness tend towards the same steady-state value as the number of neurons *N* grows sufficiently large.

However, sparseness induced by mutual inhibition is not by itself sufficient for an efficient sparse *encoding* of the input. With random receptive fields, decoding error *increases* with sparser activity.

### Stability and response to perturbations

In most machine learning experiments, the input data is randomised so that its distribution is mostly homogeneous over time. This is not the case for embodied agents that learn continuously: an animal samples from small regions of the input space as it moves from one place or activity to the next. Thus an important challenge in artificial neural networks is to learn online on non-homogeneous data. Sparse coding networks with a homeostatic term make an explicit assumption that the average firing rate of each neuron is constant, and the violation of that assumption could be a factor in catastrophic forgetting. The next experiment aims to explore whether the absence of a homeostatic term in our model makes it more robust to perturbations.

In fig. 9, we first train the network on the full MNIST dataset with Gaussian noise (*σ* = 0.2) added to the digits and clipped to [0, 1]. After 150000 stimuli, we remove the MNIST input and continue training on the background noise. We restore the input and train again on the full MNIST dataset for 150000 stimuli.

**Figure 7:**
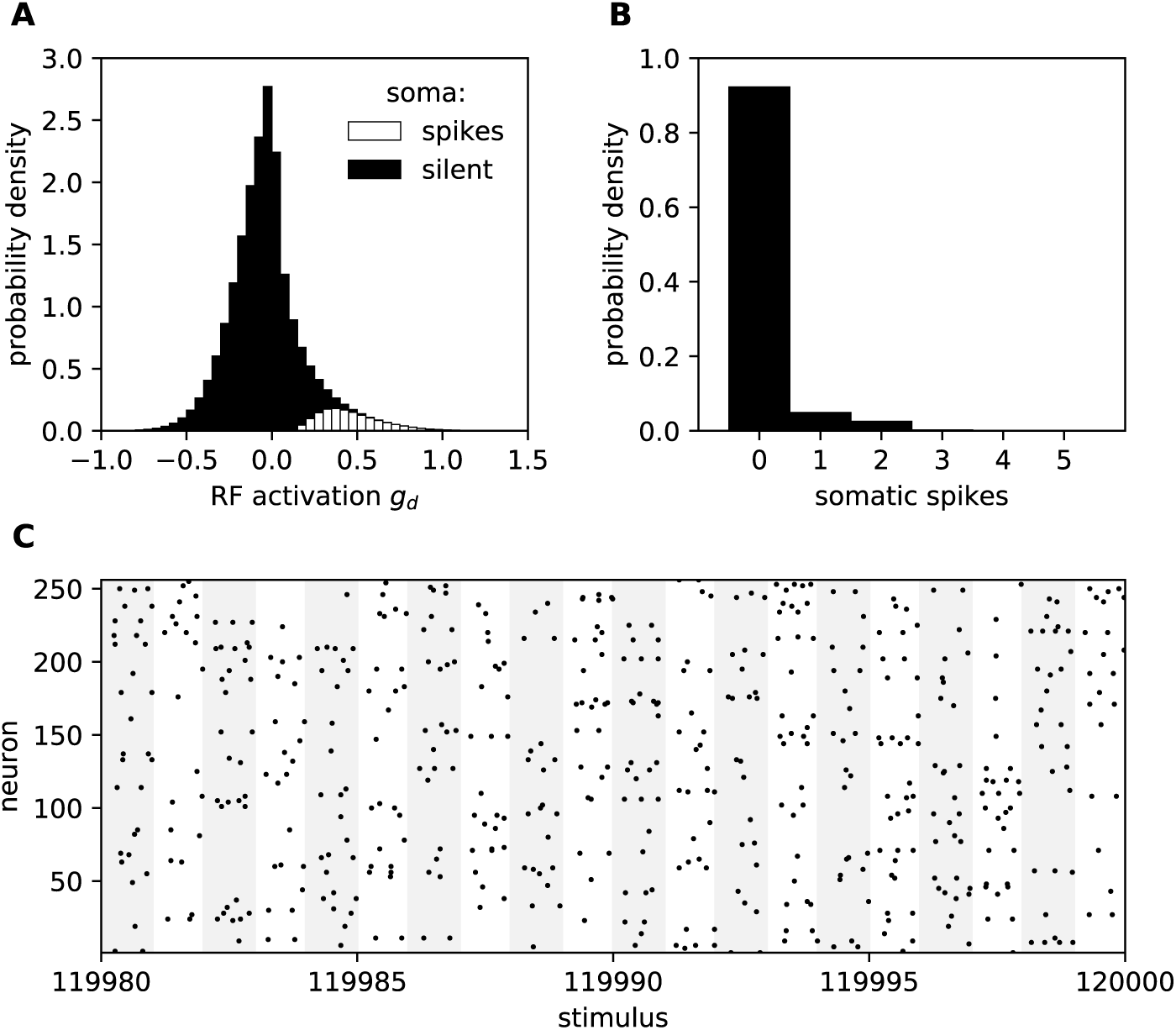
Lifetime and population activity is sparse after training on the MNIST dataset. **A:** the net dendritic input (receptive field activation) has a heavy-tailed distribution (excess kurtosis: 2.82, skewness: 0.9). The two conditions (spikes / no spikes) are stacked, not overlaid. **B:** neurons are silent most of the time, as shown by the distribution of the number of spikes per neuron per stimulus. **C:** only a few spikes are emited for each stimulus. The alternating columns in the background are 50.0 ms wide and correspond to successive MNIST patterns during a one-second period at the end of the training run.

**Figure 8:**
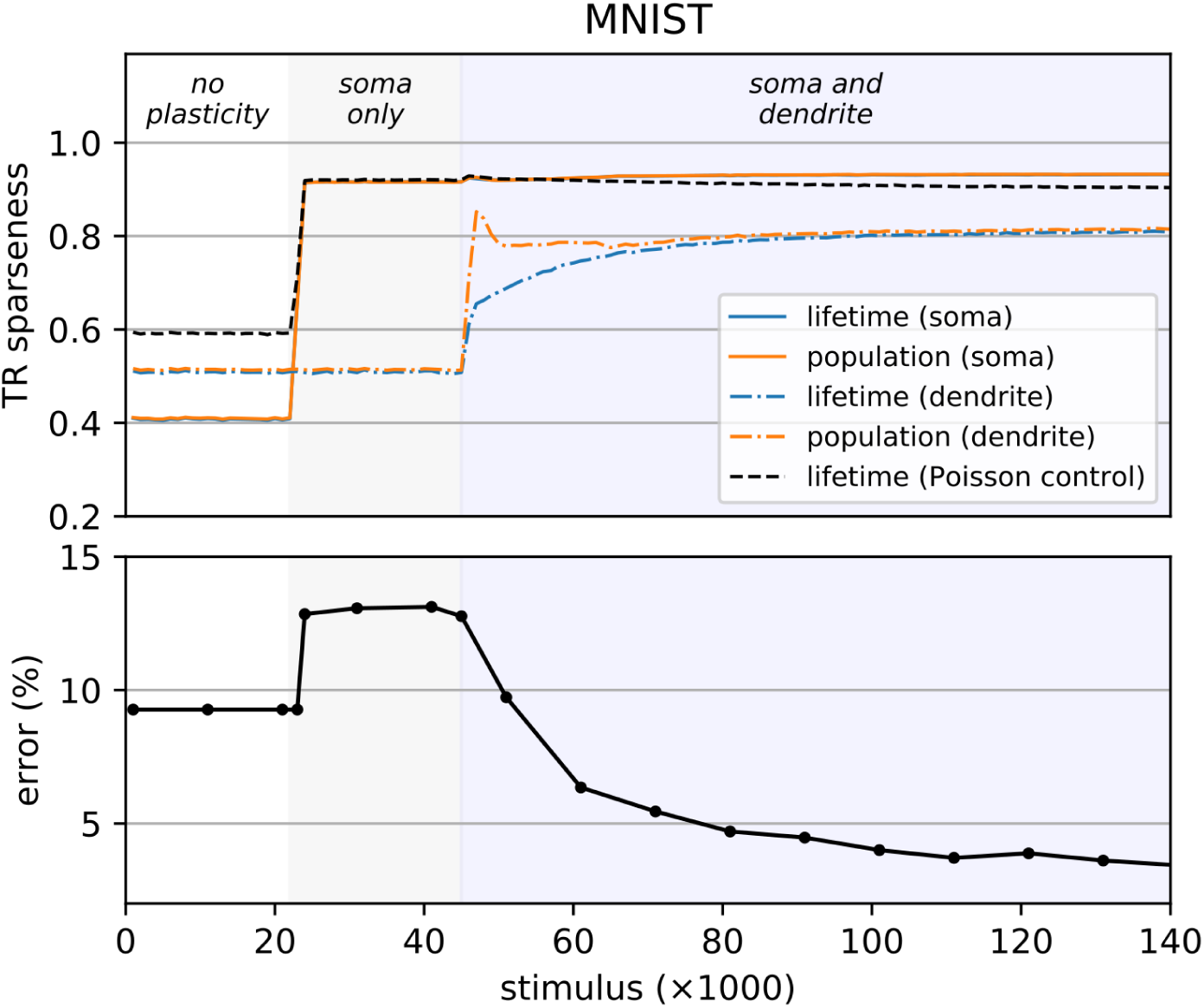
Plastic recurrent inhibition enforces sparse spiking, but feedforward plasticity is required for efficient sparse coding. Treves & Rolls (TR) sparseness (top, see *Methods* for details) and test error with a linear SVM (bottom) while training a network of *N* = 256 neurons on MNIST. Here we stage the learning in three phases: first with no plasticity (fixed initial weights); then activating the recurrent inhibition learning rule (soma only); and finally with both the recurrent and feedforward learning rules (soma and dendrite). The Poisson control shows the lifetime sparseness of a Poisson process with the same rate as the soma. The curves for the lifetime and population sparseness (soma) overlap almost perfectly in this figure. We increased the initial weights *w*, and decreased *q*, so that the initial state is less sparse. See Annex for a similar figure with natural images as the input.

Finally, we perform one last training round on a subset of MNIST that contains only the zeroes, with all other digits removed.

We find that the receptive fields retain their selectivity despite fading during the period when the network receives only background noise, and recover with minimal changes when the original input is restored (fig. 10): thanks to the lack of fast IP, input deprivation does not induce catastrophic forgetting. As long as the distribution of the independent components remains the same, there is also no drift with continued learning (compare **A** and **C** in fig. 10). In contrast, we observed a constant shifting of the receptive fields when replicating other models such as the one by Zylberberg et al. [20].

**Figure 9:**
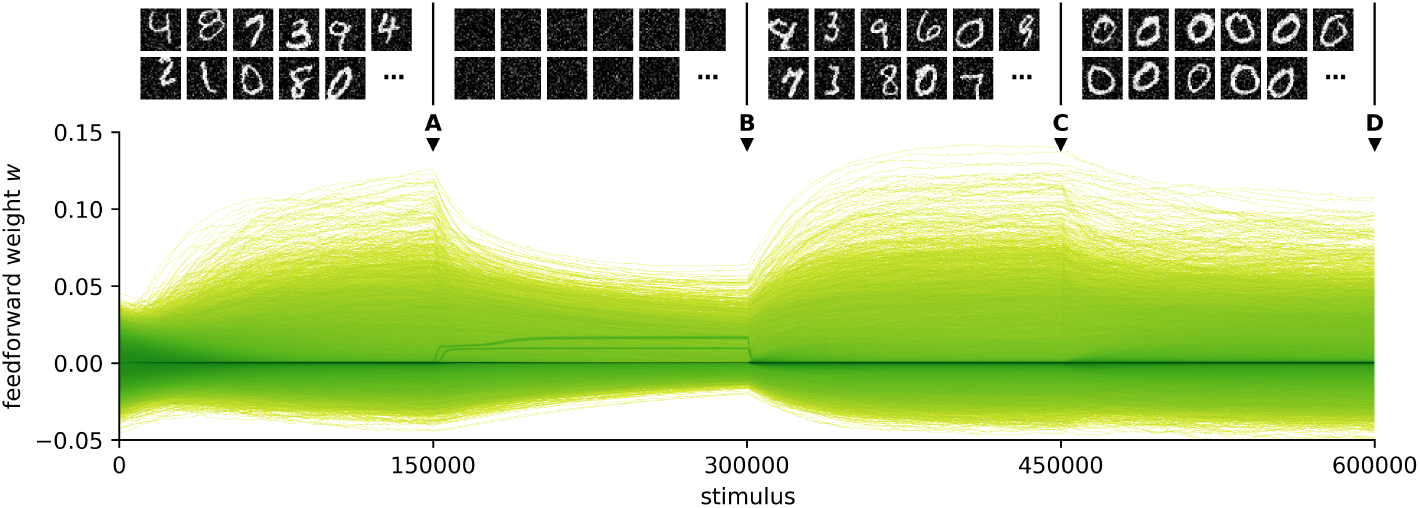
Weights converge smoothly to a steady state after perturbations in the input. **Top**: sample training stimuli for each of the four training periods. **Bottom**: density plot of the trajectories of the 112896 feedforward weights *w* during training (*N* = 144). The color mapping is logarithmic to account for the high density of zero weights. The letters **A**, **B**, **C** and **D** mark the times when snapshots of the receptive fields were taken (fig. 10).

**Figure 10:**
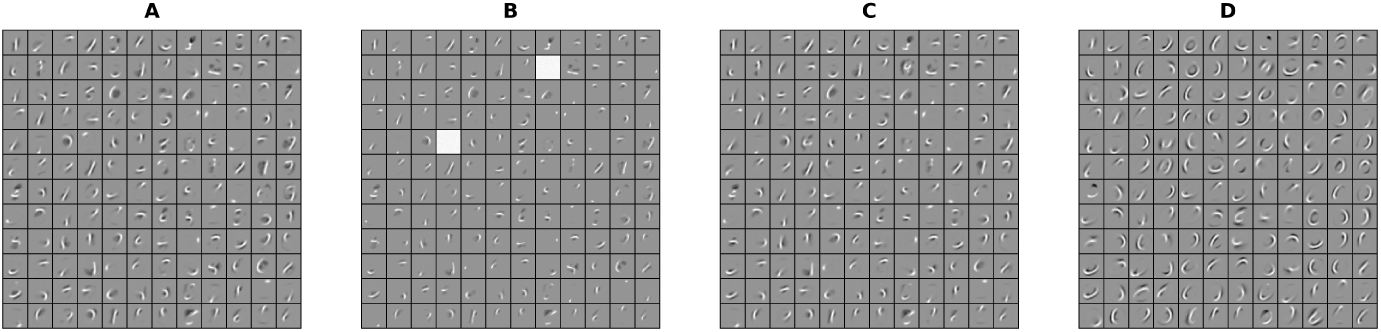
Snapshots of the receptive fields at the times marked in fig. 9. **A:** receptive fields at the end of the initial training period. **B:** receptive fields fade during input deprivation, but do not drift. **C:** they recover with minimal changes after the original input is restored. **D:** reorganisation occurs after further training with zeros only.

However, the receptive fields do change rapidly when we switch from the full MNIST to zeroes only: they adapt to match the new distribution of the independent components and forget the features that were specific to other digits (such as straight lines). Thus the lack of IP protects against forgetting during input deprivation but does not block continual adaptation to the input, as long as the new stimuli overlap with existing receptive fields.

A small number of neurons (typically one or two) respond strongly to the noise during the period of input deprivation (bright receptive fields in fig. 10; dark lines in figs. 9, 11). Average firing rates for the other cells are low; again, this can be explained by the absence of a homeostatic term that would drive every neuron towards a target firing rate. Since the background noise does not contain any structure, these few active cells are sufficient to encode it and inhibit other neurons, protecting their receptive fields.

The transient increase in activity when the input is restored does not exceed three times the baseline: spikes remain sparse throughout, and come back to normal after 10 seconds (fig. 11). Since the neurons have fixed somatic and dendritic thresholds, that increase must come from the decay of lateral inhibition or from a shift in the excitatory/inhibitory balance of the feedforward weights. In contrast, in a network with IP, homeostatic adjustment of the thresholds to the background noise would cause a temporary saturation of the transfer function and loss of sparseness when the input is restored.

**Figure 11:**
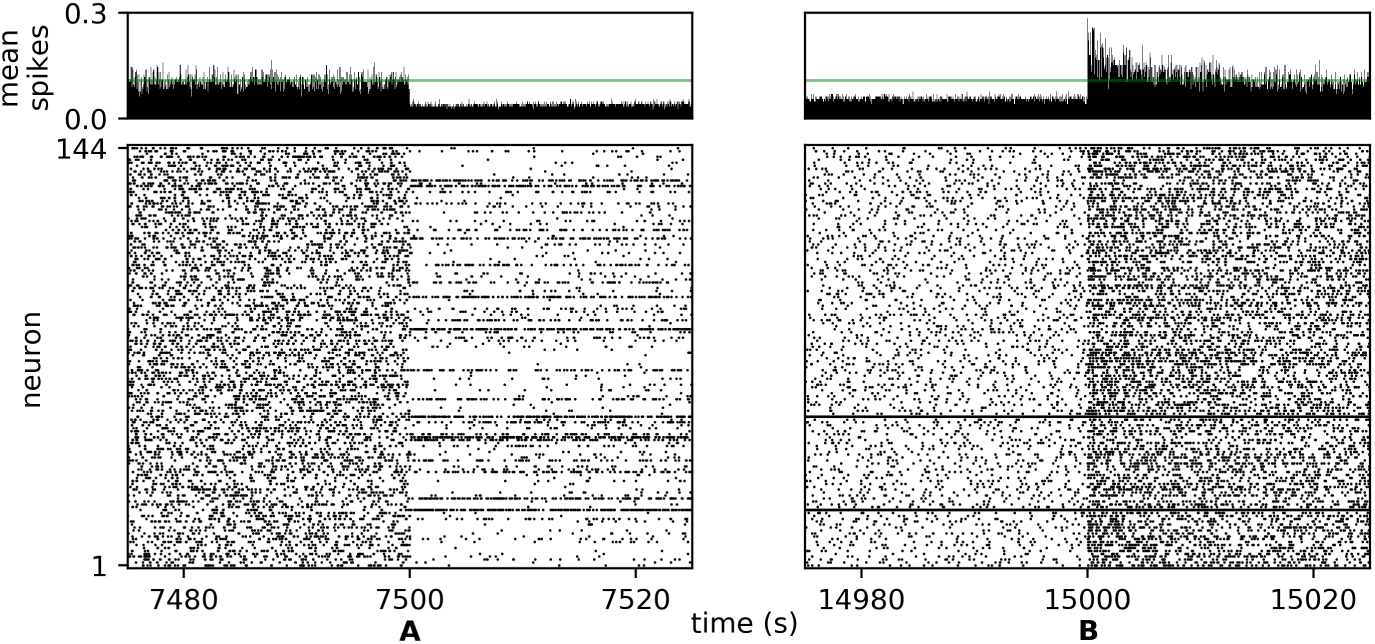
The network is robust to input deprivation. **Top:** mean number of spikes per neuron per stimulus; the green line marks the average value before mark A. **Bottom:** raster plot of the output spikes. Following input deprivation (mark **A**), firing patterns return to normal within 20 seconds after the input is restored (mark **B**), except for the neurons that responded strongly to the noise (these take somewhat longer).

## Discussion

### Sparse coding does not require fast IP

Our findings confirm that intrinsic plasticity (IP) is not strictly required to learn sparse codes. Plastic lateral inhibition, in addition to its role in decorrelating the population responses, can also regulate sparseness through its effect on the nonlinear Hebbian learning rule.

The idea of enforcing population sparseness is not new — in networks where learning uses global information, one can devise a cost function with a sparseness constraint based on population activity, and minimise it to learn a suitable set of receptive fields.

But this global cost information is not typically available in neural networks that use only local information for learning, in line with biology. Thus previous models [15,19,20] have made use of IP to enforce lifetime sparseness, as this information is readily available in each neuron. We show that this is not the only way: information about population sparseness *can* be conveyed to local rules through mutual inhibition, extracted thanks to input compartmentalisation, and used to learn independent components in the same way as lifetime sparseness.

When it comes to explaining how cortical neurons might learn sparse codes, this helps to resolve a conflict of timescales. The mechanism enforcing sparseness must be faster than Hebbian learning to avoid unstability, and consequently IP is fast in models that rely on it for sparse coding. But this conflicts with what we know about IP in biological neurons: homeostatic firing rate adaptation is normally quite slow, on the order of hours or days [28–30].

Enforcing sparseness via lateral inhibition instead is a better match for the data: fast adaptation of these connections is plausible through a combination of short-term facilitation and long-term potentiation of inhibitory synapses, which can occur over a timescale of seconds to minutes. Freed from the task to stabilise Hebbian learning, IP could have other computational roles on slower timescales, for instance helping to recruit previously silent neurons and dendrites or adapting to ongoing slow processes like structural plasticity and developmental changes.

### Compartmentalised inputs let local rules estimate population sparseness

Our learning rule has access to information about population sparseness thanks to the separation of the feedforward and recurrent pathways. If these were integrated together into a single activity variable, there would be no way to distinguish weaker inputs from stronger competition. Compartmentalised integration, with feedforward activity integrated in the dendrite and lateral inhibition integrated in the soma, disambiguates the reasons why a neuron does not fire and allows each neuron to compute a local estimate of the amount of competition with its neighbours.

Point neurons have been cost-effective approximations wherever simulation of thousands of neurons in real time is the aim, and moving away from that paradigm requires good reasons. Our model gives another example of the types of learning that become possible in neurons with compartmentalised inputs and may justify the expense. In contrast to multi-compartmental models with detailed branching and morphology, neurons with a few input compartments are not much more expensive to simulate than point neurons, requiring only a handful of extra state variables. This makes them suitable for large-scale simulations and hardware implementations — the SpiNNaker and Loihi neuromorphic chips, for instance, already support compartmentalised inputs [42,43].

### Sparse coding via population sparseness is robust to input deprivation

Without fast IP, the network can be made more robust to temporary input deprivation. If the input is replaced by background noise, lateral inhibition will adjust in seconds, just as fast IP would rapidly adjust the firing threshold. But the important information is in the receptive fields of the dendrites, and in our model these do not adapt rapidly when the neurons are exposed to noise.

Dendrites respond weakly to noise because noise, lacking structure, tends to activate their excitatory and inhibitory receptive fields equally. At the population level, a couple of neurons will eventually develop broad excitatory receptive fields and start responding to the noise. But because there is no structure in noise to support a division of labour between output neurons, the first few responders will be able to inhibit all the others. This keeps the overall activity low and protects most receptive fields from change: the feedforward learning rule (eq. 3) is gated by post-synaptic activity and weights will not change fast if the dendrite is weakly active and the soma is inhibited.

In contrast, in models with IP, input deprivation causes a rapid adjustment of the spiking or plasticity thresholds to maintain the same average firing rate. Consequently the rate of feedforward synaptic changes stays high, and receptive fields are lost to the background noise.

But replacing intrinsic plasticity with synaptic plasticity is still not enough to cope with other types of changes, such as those that an animal or robot would encounter as it switches between tasks and environments: the network remains susceptible to rapid and extensive reorganisation when novel inputs overlap the existing receptive fields, or when the distribution of the independent components changes. On the one hand, that kind of adaptability is desirable as natural environments are not static and the quick acquisition of novel stimuli can be critical for survival. On the other hand, it should disturb existing receptive fields as little as possible so as not to erase previous experiences and all the associations that build upon them.

Although increasing sparseness and careful tuning of learning rates could help, it is likely that solving that stability-plasticity dilemma will require ad-hoc gating mechanisms. Some candidates are the conditional consolidation of synaptic changes [44], neuromodulation and attention [45,46], or a mechanism based on top-down prediction errors like the Adaptive Resonance Theory [47].

### Sparse activity does not imply sparse coding

We observe a dissociation between measures of sparseness and measures of decoding accuracy: high sparseness is achieved almost immediately via the potentiation of lateral inhibitory weights, while high decoding accuracy requires adequate receptive fields learned by the feedforward plasticity rule. In fact decoding accuracy in a network with random receptive fields decreases with sparseness. Conversely Zylberberg & DeWeese [48] observed decreasing sparseness as a result of receptive field formation in their sparse coding network.

Thus there is more to sparse coding than just sparse activity. Sparseness is a constraint for learning, not by itself a guarantee of an efficient encoding: sparse coding requires receptive fields that match the independent components of the input, so that the same amount of information can be transmitted with fewer spikes and fewer active units.

Lehky et al. [41] caution that high measures of sparseness can be obtained trivially via non-linear, information-discarding transforms. For instance the average lifetime sparseness could be increased simply by raising the thresholds of the neurons, without learning selective receptive fields; but this would also decrease the information transmitted about the input. They argue that statistical measures of sparseness are not a sufficient optimality criterion for explaining the architecture of nervous systems, and that information transmission and other ecologically-relevant aspects must also be considered.

Here we evaluate linear decodability, in the light of the hypothesis that the computational cost of decoding is a major biological constraint together with coding density and metabolic efficiency, and that linear decoding requires fewer synapses. Other criteria could be also be examined, such as whether output correlations are still relevant for decoding [49], or robustness to noise and to the loss of coding units.

### Comparison with similar models

Our model is far from being the first to learn features similar to the ones observed in V1 cells, but it differs from previous models in several ways. The learning rule used by Olshausen & Field [1] minimises the output reconstruction error, as do sparse auto-encoders [8], whereas our model does not need or compute that information. This reduces the number of steps required for learning, and as a model of sparse coding in the brain it requires neither multiple layers nor a biological mechanism for the backpropagation of error. Földiák [15], Zylberberg et al. [20] and King et al. [21], and models based on the standard BCM rule rely on homeostatic threshold adaptation, while Savin et al. [19] also use IP to adjust the transfer function of the neuron. Our model uses fixed thresholds instead, both in the dendrite and the soma, relying on population rather than lifetime sparseness as discussed before.

The term *x*(*z* − *δy*) at the core of the feedforward learning rule is reminiscent of the error-correcting Delta rule [50,51], and it can be seen as a rate-based relative of the rules used in Körding & König [31] and Urbanczik & Senn [32].

But in Urbanczik & Senn, the purpose of learning is to correct the mismatch between the somatic activity, *z*, and its prediction by the dendrite, *φ*(*y*), so that the dendrite learns to move the somatic membrane potential towards the equilibrium potential of predictable somatic inputs. After learning, that dendritic prediction is mostly correct, *z* ≈ *φ*(*y*) and the predicted somatic inputs have no effect.

In contrast, the goal of our learning rule is not to achieve a perfect prediction of the somatic inputs by the dendrite, but to exploit the mismatch between *z* and *y* so that it creates a BCM-like curve modulated by inhibition. Thus the steady state does not occur at *z* ≈ *δy*. Starting from the learning rule: d*w* = *x*(*z* − *δy*) − *yw*, let us treat *x*, *w*, *z* and *y* as correlated random variables. Then, for 〈d*w*〉 = 0, we have 〈*x*(*z* − *δy*)〉 = 〈*yw*〉. Substituting *w* with a constant *w*_∞_, we get the following non-trivial fixed point:

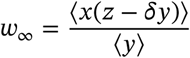

In other words, the fixed point over a certain set of inputs is when each weight equals the mean of the loser / winner function *z* − *δy* (fig. 2) times the input, normalised by the mean dendritic activity. This yields receptive fields which are inhibitory for input dimensions associated with losing the competition (*z* < *δy*) and excitatory for those associated with winning (*z* > *δy*). If there was no mismatch between *z* and *δy* in the steady state, the receptive fields would be blank (*w*_∞_ = 0).

The model by Körding & König [31] is perhaps the closest to our work. In their network, lateral inhibition decides whether a neuron is losing or winning the competition by blocking the backpropagating action potentials, switching the sign of plasticity in the dendrite. However, it does so without blocking the spikes that travel down the axon, whereas lateral inhibition in our model suppresses both the internal teaching signal and the output of the neurons. The distinction could have its relevance in multi-layer networks; but it is likely that both architectures can perform sparse coding with the right dendritic learning rule — something that Körding & König [31] did not explore, as they used simple stimuli such as moving bars which do not contain multiple independent components.

### Biological interpretation

Our model is only loosely based on biology: at its core, it is mainly a computational exploration of compartmentalised input integration in the context of sparse coding, and whether biological neurons make use of similar principles remains an open question. But it does suggest phenomenological interpretations for a number of experimental facts.

In terms of architecture, there are multiple inhibitory pathways in the cortex [52], and some of these pathways target dendrites or somas specifically as they do in our model; for instance the fast-spiking, parvalbumin-positive basket cells mediate lateral inhibition preferentially via synapses close to the soma. There are, however, many other pathways, including recurrent inhibitory pathways targetting dendrites, that our model does not explain.

As for synaptic plasticity, let us rearrange the terms of the feedforward learning rule (eq. 3) and annotate it to match the terminology used in neuroscience:

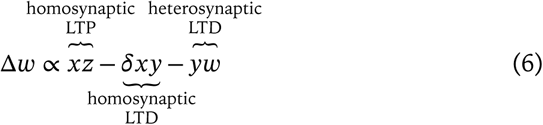

where homosynaptic refers to plasticity induced in the synapse that was stimulated (correlated pre and post activity), and heterosynaptic refers to plasticity induced in other synapses (independently of whether they were active).

This highlights a number of testable hypotheses. First, it assumes that homosynaptic LTP (*xz*) is induced by correlated preand post-synaptic spikes, which is well established by a long history of electrophysiological experiments from Hebb [2] to STDP theory [53].

Heterosynaptic LTD (−*yw*) is attested in some neurons [54,55] as a form of normalisation of total synaptic input and may be linked to competition for metabolic resources between synapses. While developing our model we tested a variant (−*zw*) for this term which depended *z* on instead of *y*. This yielded inferior decoding performance but similar receptive fields, therefore we do not want to make a strong claim of the dendritic vs. somatic dependence of heterosynaptic LTD.

Less obvious is the notion that correlated *dendritic* activity should induce homosynaptic LTD (−*xy*): We know, on the contrary, that dendritic spikes can sometimes induce LTP on their own [56,57]. But we also know that NMDA receptor activation can induce LTD or act as a negative feedback on potentiation [57,58], which is compatible with our learning rule.

In terms of learning paradigms, our model makes hypotheses that diverge from the classical framework of spike timing-dependent plasticity [53], where low firing rates tend to induce homosynaptic LTD and high firing rates tend to induce homosynaptic LTP in a way that is compatible with the standard BCM rule [59].

In contrast, our model assumes that low post-synaptic rates cause homosynaptic LTD of dendritic synapses only when they are due to strong recurrent inhibition, and not to a weak feedforward input. Conversely, it assumes that recurrent inhibition can switch the sign of plasticity at dendritic synapses and turn LTP into LTD, an idea that has been suggested in computational models [31,60] but requires further experimental validation.

Other aspects of biological neurons, however, are more difficult to reconcile. For instance, our model relies on lateral inhibition shifting the response curve of the somas to the right – a phenomenon known as subtractive inhibition, in contrast with divisive inhibition which modulates the response slope without changing its threshold. But in pyramidal neurons somatic inhibition is normally divisive rather than subtractive: it is dendritic inhibition that has a subtractive effect [61]. It may be that the mechanisms we distribute over a somatic and a dendritic compartment, occur *within dendrites* in biology, possibly involving compartmentalisation between the dendritic shaft and the spines, or between different dendritic variables like voltage and calcium.

More generally, interpreting our learning rule calls for further electrophysiological investigations of how different pathways contribute to synaptic plasticity — looking not just at pairs or triplets of pre and post spikes, but also at coincident dendritic activity and inhibitory modulation.

### Relevance for machine learning

Most of the recent advances in machine learning have relied on supervised learning, and there have been efforts to make supervised learning algorithms like error backpropagation work with spiking neurons as well [62,63]. But there is also a notion that unsupervised learning will play an increasing role in future learning systems. A spiking neural network learning sparse codes with local, unsupervised rules, and running on neuromorphic hardware, would have several advantages over current approaches that use dense networks of rate neurons. It could use considerably less energy due to an efficient matching of sparse activity with sparse, event-based communication. It could replace the first layers of deep neural networks, which tend to learn the same sort of features, but it could also form the basis for new hierarchical architectures like the ones proposed by Hawkins & Ahmad [64]. And unlike batch training algorithms it would be suitable for online learning that adapts continuously to new data.

## Models & Methods

### Somatic compartments and somatic synapses

The somatic compartments are standard LIF neurons. The membrane potential *u* follows the following equation:

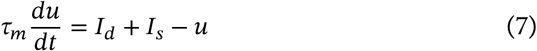

where *I*_*d*_ and *I*_*s*_ are the currents from the dendrite and somatic synapses, respectively.

We use a fixed spiking threshold *θ* and after-spike reset *ρ* without a refractory period:

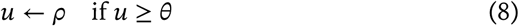

We compute a firing rate *z* that takes into account the number of spikes and also their latency relative to the stimulus onset *t*_0_, with the aim of producing a smooth measure that is sensitive to small changes in activity even in the case of a single spike. First we define a trace ζ that increases after each spike and decays exponentially:

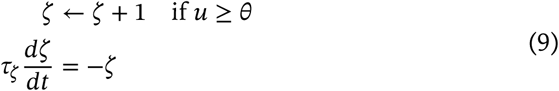

Then we normalise so that the area under the curve is the number of spikes, and integrate over the stimulus window:

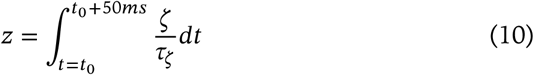

Thus a spike that occurs towards the end of the window contributes less to the total than a spike that occurs early. This also approximates the effect of input eligibility traces in more detailed models.

Somatic inhibition comes from lumped, conductance-based synapses where the fraction *g*_*s*_ of active conductance and somatic current *I*_*s*_ evolve as follows for each neuron post:

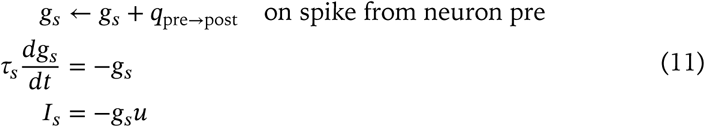

The initial weights *q* are drawn from an exponential distribution (mean = 0.01). Before each new stimulus we reset the continuous-time variables of the model, as in Zylberberg et al. [20]:

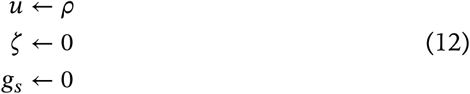

That reset does not seem to be critical for our findings, but we did not explore the issue further.

### Dendritic compartments

Dendrites are rate- and current-based. The net dendritic input *g*_*d*_ and dendritic activation _*y*_ for each neuron post are as follows:

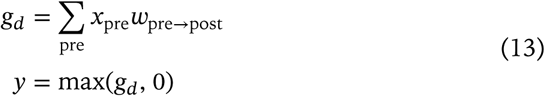

The initial weights *w* are drawn from a normal distribution (std = 0.01).

The current *I*_*d*_ from the dendrite to the soma is a nonlinear function of the dendritic activation:

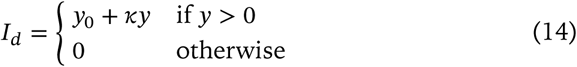

Here the goal is to reproduce the active properties of biological dendrites. Above a certain input threshold, regenerative activation of the NMDA receptors causes dendritic spikes. These lead to a sharp increase in membrane potential followed by a plateau where stronger inputs cause no further increase in voltage [65,66]. We model this with a step function and the offset *y_0_*. However, stronger inputs do increase the duration, and reduce the rise time of the plateau, producing more somatic spikes. We model this with the linear term *κy*. In practice we adjust *y*_0_ to cancel out the somatic rheobase, so that suprathreshold dendritic activation elicits at least one spike in the absence of somatic inhibition. Dendrites start responding when *g*_*d*_ > 0. We find that the actual threshold is not critical for our findings as long as all dendrites respond to some inputs at the start of the simulation. A small positive value would better reproduce the data in Milojkovic et al. [65] and Oikonomou et al. [66].

The coupling between the soma and the dendrite is one-way: somatic potentials have no effect on dendritic activity in our model. This ignores the effect of backpropagating action potentials on dendritic voltage, but it does match the data on somatic inhibition, which has almost no impact on the generation of dendritic spikes [67,68].

### Measure of sparseness

In fig. 8 we use the same measure of sparseness as Zylberberg & DeWeese [48], called TR sparseness after Treves & Rolls and related to the coefficient of variation. It is computed as follows:

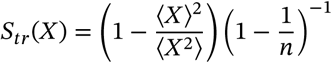

where *n* is the length of *X*.

For lifetime sparseness, *X* is a vector of 1000 observations from the same neuron over time, and we then average *S*_*tr*_ across all neurons in the population. For population sparseness, it is a vector of observations from all *N* neurons across the population at a particular instant, and *S*_*tr*_ is averaged over 1000 stimuli. Somatic sparseness uses the number of spikes (one could use the output rate *z*, but the number of spikes is easier to compare to a Poisson control) while dendritic sparseness uses the dendritic activity *y*.

In our case the Gini index gives qualitatively similar results to TR sparseness, and could be used interchangeably. Both are more stable over time than excess kurtosis, as noted by Hurley & Rickard [69].

### Parameters

Table 3 summarises the parameters for the neuron model. All simulations are performed with a timestep *dt* = 0.5 ms and 100 steps per stimulus, except for fig. 2 which uses a finer timestep.

**Table 3:** Model parameters used for all experiments in this paper.

| Soma and somatic synapses | Dendrites |
| --- | --- |
| $\theta = 1.0$ | $y_0 = 1.0$ |
| $\rho = 0.0$ | $\kappa = 0.5$ |
| $\tau_m = 10.0$ ms | $\delta = 0.5$ |
| $\tau_\zeta = 50.0$ ms | $\lambda = 0.01$ |
| $\tau_s = 5.0$ ms | $\mu = 4 \times 10^{-4}$ |
| $\nu = 1 \times 10^{-1}$ | |
| $\beta = N/250$ | |

### Receptive Fields

Throughout this paper we use the weights of the neurons as a proxy for their actual receptive fields. Showing all the weights of the network on the same image requires that we normalise each receptive field separately, because neurons that respond to narrow features have larger absolute weights than those that respond to broad ones. Nonetheless, we make sure that zero weights appear as the same middle gray for all neurons, allowing quick identification of ON (brighter) and OFF (darker) areas. Thus we normalise the receptive field *W*_*i*_ = [*w*_1→*i*_ … *w*_*d*→*i*_] of each neuron *i* as follows when generating the figures:

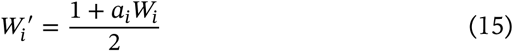

where *a*_*i*_ = (max_1≤*j*≤*d*_|*w*_*j*→*i*_|)^−1^ and *d* is the number of input dimensions.

### MNIST

We use both the standard MNIST dataset [34] and the Fashion-MNIST variant [35], each with 60,000 training samples and 10,000 test samples. We map the full range of the data to the interval. When training the sparse coding network, we shuffle the patterns and distort them with random shears and translations, as done in LeCun et al. [39]. The purpose of these distortions is to increase the number of distinct training samples, and also to remove the correlations introduced by the centering of the patterns. We do this by applying the following affine transformation with the origin at the center of the pattern:

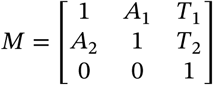

where each *A_i_* is a random variable drawn from *N*(σ = 0.1), and each *T*_i_ is a random variable drawn from *N*(σ = 2.0). Distorted digits produce more localised receptive fields than the centered patterns, which in turn improves the receptive fields than the centered patterns, which in turn improves the performance of classifiers trained on the output of the network. When training and testing the classifiers themselves, we freeze the weights of the sparse coding network and we use the plain stimuli without distortions.

In tbl. 1 we use the following classifiers from `scikit-learn` [70] version 0.19.1:

**SVM:** `LinearSVC(C=1.0, class_weight=None, dual=False, fit_intercept=True, intercept_scaling=1, loss=squared_hinge, max_iter=1000, multi_class=ovr, penalty=l2, random_state=2136146589, tol=0.0001, verbose=0)`

**kNN:** `KNeighborsClassifier(algorithm=auto, leaf_size=30, metric=minkowski, metric_params=None, n_jobs=1, n_neighbors=4, p=2, weights=uniform)`

### Natural images

We use two datasets of photographic images: one by Olshausen & Field [1], and one compiled from public-domain archive images of monuments from the Cornell University Digital Collections [36]. In both cases, each image was converted to grayscale, resized to an area of 200,000 pixels, preprocessed using the same whitening transform as Olshausen & Field [1], and then normalised to unit variance. No further normalisation was applied to the individual patches used for training; in particular, the patch mean was not subtracted from the input. Note that in contrast to MNIST the natural image stimuli contain both positive and negative values. We interpret these as ON and OFF channels from the retina; while it would be more realistic to split the ON and OFF values into separate, non-negative channels, we did not attempt this here.

For the reconstruction experiment, the input image was tiled into overlapping patches with a width of 16 pixels and a stride of 8 pixels. Each input patch was run through a sparse coding network pre-trained on the Monuments dataset. The sparse output was then fed as the input to a linear model trained with ridge regression to recontruct the original patches. Finally, the predicted patches were placed at their original locations and averaged to account for the stride overlap.

## Acknowledgements

DD and VVH were funded by the GRK 1589/1 and 1589/2 Sensory Computation in Neural Systems of the Deutsche Forschungsgemeinschaft (DFG).

DD and VVH were funded by EU FP 7 Grant 609465 (EARS).

DD and MS were funded from EU FP 7 Grant 604102 and H2020 Grants 720270 and 785907 (Human Brain Project SGA1 and SGA2).

## Annex

**Figure 12:**
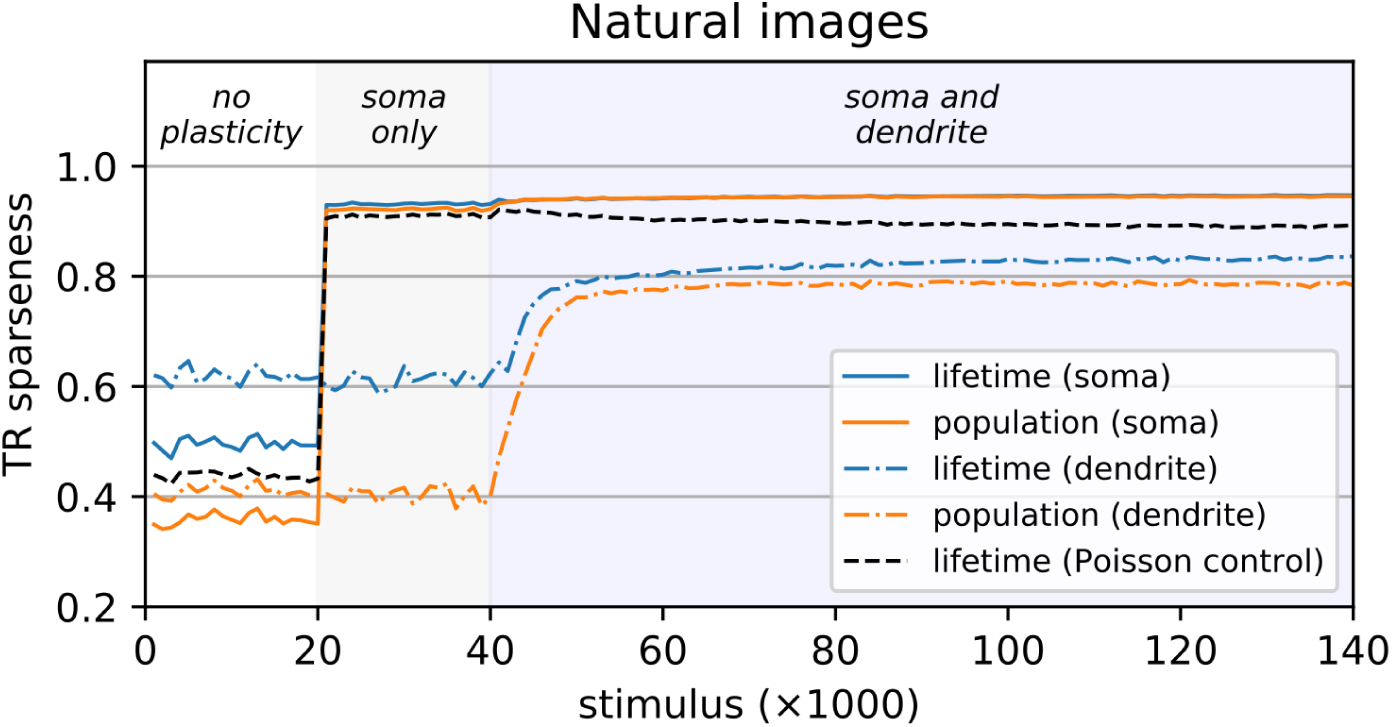
Sparseness follows a similar timecourse with natural images. Same as the top part of fig. 8, but this time training on the Monuments dataset of natural images instead of MNIST. In this case we did not compute a reconstruction error.

